# Role for the flagellum attachment zone in the resolution of cell membrane morphogenesis during *Leishmania* cell division

**DOI:** 10.1101/2020.03.26.009928

**Authors:** Clare Halliday, Ryuji Yanase, Carolina Moura Costa Catta-Preta, Flavia Moreira-Leite, Jitka Myskova, Katerina Pruzinova, Petr Volf, Jeremy C. Mottram, Jack D. Sunter

## Abstract

The shape and form of the flagellated eukaryotic parasite *Leishmania* is sculpted to its ecological niches and needs to be transmitted to each generation with great fidelity. The shape of the *Leishmania* cell is defined by the sub-pellicular microtubule array and the positioning of the nucleus, kinetoplast and the flagellum within this array. The flagellum emerges from the anterior end of the cell body through an invagination of the cell body membrane called the flagellar pocket. Within the flagellar pocket the flagellum is laterally attached to the side of the flagellar pocket by a cytoskeletal structure called the flagellum attachment zone (FAZ). During the cell cycle single copy organelles duplicate with a new flagellum assembling alongside the old flagellum and these are then segregated between the two daughter cells by cytokinesis, which initiates at the anterior cell tip. Here, we have investigated the role of the FAZ in the morphogenetic resolution of the anterior cell tip during cell division. We have deleted the FAZ filament protein, FAZ2 and investigated its function using light and electron microscopy and infection studies. The loss of FAZ2 caused a disruption in membrane organisation at the anterior cell tip, resulting in cells that late in division were connected to each other by a membranous bridge structure between their flagella. These changes had a great impact in vivo with the FAZ2 null mutant unable to develop and proliferate in sand flies and causing a reduced parasite burden in mice. Our study provides a deeper understanding of membrane-cytoskeletal interactions that define the shape and form of an individual cell and the remodelling of that form during cell division.

**Author summary:** *Leishmania* are parasites that cause leishmaniasis in humans with symptoms ranging from mild cutaneous lesions to severe visceral disease. The life cycle of these parasites alternates between the human host and the sand fly vector, with distinct forms in both. These different forms have different cell shapes that are adapted for survival in these different environments. *Leishmania* parasites have an elongated cell shape with a flagellum extending from one end and this shape is due to a protein skeleton beneath the cell membrane. This skeleton is made up of different units one of which is called the flagellum attachment zone (FAZ), that connects the flagellum to the cell body. We have found that one of the proteins in the FAZ called FAZ2 is important for generating the shape of the cell at the point where the flagellum exits the cell. When we deleted FAZ2 we found that the cell membrane at the tip of the was distorted resulting in unusual connections between the flagella of different cells. We found that the disruption to the cell shape reduces the ability of the parasite to infect mice and develop in the sand fly, which shows the importance of the parasite shape.

## Introduction

The kinetoplastid parasites have a defined shape and form, which varies during their life cycle depending on the specific ecological niche. These different shapes and forms of *Leishmania spp* and *Trypanosoma brucei* are determined by their highly organised sub-pellicular microtubule array. Different forms can be categorised based on the relative positions of three key structures:- i) the nucleus, ii) the kinetoplast (the condensed mitochondrial DNA) and iii) the flagellum and its associated flagellar pocket (an invagination of the cell membrane at the base of the flagellum) (1). The flagellar pocket is a critical feature as it is the site of all endo- and exocytosis in these parasites.

The *Leishmania* flagellar pocket consists of two regions, the bulbous lumen and the flagellar pocket neck. At the distal end of the bulbous lumen is the flagellar pocket collar, a cytoskeletal structure that cinches in the cell membrane to form this bulbous domain (2). Distal to this point, within the flagellar pocket neck region, the cell body membrane is closely apposed to the flagellum membrane until the flagellum exits the cell body. Since the *Leishmania* promastigote flagellum extends from the anterior cell tip it has traditionally been described as ‘free’; however, the basal region is firmly attached to the cell body within the flagellar pocket neck region (2).

The attachment of the flagellum to the cell body is mediated by the flagellum attachment zone (FAZ), a complex structure, which contains cytoskeletal elements (3,4). The FAZ connects the cell body cytoskeleton to the cytoskeleton of the flagellum through the cell body and flagellum membrane and consists of three major domains:- i) the cell body domain, ii) the intermembrane domain, and iii) the flagellum domain (4). The *T. brucei* FAZ is much longer than the *Leishmania* equivalent and has been the subject of more extensive study. In *T. brucei* electron microscopy studies have defined the individual cytoskeletal elements that together form the FAZ. Within the cell body domain there is the FAZ filament that runs parallel to the microtubule quartet (a specialised set of four microtubules) that together form a seam within the sub-pellicular microtubule array (4,5). From the FAZ filament and microtubule quartet a set of fibres extend to a linear array of regular junctional complexes embedded in the cell body membrane. These junctional complexes connect across to the flagellum membrane and together with that membrane form the intermembrane domain. Another set of intraflagellar filaments then constitute the linker structure that connects to the proximal domain of the paraflagellar rod (an extra-axonemal structure) (6).

Work in many labs and more recently the genome-wide tagging project, TrypTag has identified many FAZ proteins in *T. brucei* (7–12). The function and interaction of a number of these proteins has been investigated; for example, the depletion of FAZ2 resulted in full length flagellum detachment and the loss of other FAZ proteins such as FAZ1 and FAZ8 (12).

The *Leishmania* orthologs of many *T. brucei* FAZ proteins localised to the *Leishmania* flagellar pocket neck region, where the FAZ in *Leishmania* is found (2). Despite the conservation in protein content there are two major organisational differences between the cytoskeletal elements of the *Leishmania* and *T. brucei* FAZ (2). Firstly, in *Leishmania* the microtubule quartet and FAZ filament are not found beneath the primary region of flagellum-to-cell body attachment but are positioned approximately a quarter turn anti-clockwise from this attachment region, in cross sections through the FAZ area looking towards the posterior of the cell (2); however, near the anterior cell tip the attachment region broadens and here the distal end of the microtubule quartet and FAZ filament connect into the attachment zone proper. Secondly, in *Leishmania* the fibres of the FAZ within the flagellum do not connect with the PFR but instead connect the flagellum membrane with the axonemal microtubule doublets (2). Despite the physical separation between the cytoplasmic components of the FAZ and the primary attachment region in *Leishmania*, there is still a clear connection between the flagellum and the cell body membranes in this region, and the membrane attachment area is linked to the flagellar cytoskeleton.

In *T. brucei* the FAZ has additional functions beyond maintaining lateral flagellum attachment to the cell body, including a key role in cytokinesis and cell morphogenesis. During the cell cycle a new flagellum and associated FAZ are assembled alongside the existing flagellum and the distal end of the FAZ is the site for cytokinesis furrow ingression (13,14). Moreover, the distal end of the growing FAZ is associated with a complex of proteins that are important for cytokinesis (McAllaster *et al.*, 2015; Zhou *et al.*, 2016a; Zhou *et al.*, 2016b; Kurasawa *et al.*, 2018; Zhou *et al.*, 2018; Hu *et al.*, 2019) and, the depletion of FAZ proteins such as ClpGM6 and FLAM3, reduced FAZ length, resulting in the misplacement of the cleavage furrow and the production of shorter cells (21,22).

We have recently begun to investigate the function of the FAZ in *Leishmania* and have interrogated the function of FAZ5, a protein with multiple transmembrane domains that is likely a component of the primary attachment region (23). The deletion of FAZ5 caused the loss of flagellum attachment to the cell body along the flagellar pocket neck region but this did not affect the growth of these cells in culture and there were no obvious cytokinesis defects. This contrasts to *T. brucei,* where flagellum detachment by depletion of FAZ proteins such as FLA1, FAZ2 and CC2D caused cytokinesis defects and cell death (8,10,12). However, the FAZ5 mutant cells were shorter and wider, with an altered flagellar pocket architecture, indicating that the FAZ in *Leishmania* has a role in cell morphogenesis (23).

The separation of the primary attachment region from the FAZ filament and microtubule quartet in *Leishmania* provided an opportunity to interrogate the function of these parts in isolation from the other FAZ domains (2). Here, we focussed on the function of FAZ2 in *Leishmania* as in *T. brucei* this protein was identified as an essential FAZ filament component (12). Deletion of FAZ2 in *Leishmania* was not lethal; however, many cells displayed a distinct phenotype: cells late in division were connected to each other by a membranous bridge structure between their flagella. Thus, although not a prominent feature of *Leishmania* the phenotype described here reveals an important role for the FAZ filament in cell segregation and morphogenesis.

## Results

### Deletion of FAZ2 impaired cell segregation after cytokinesis

To understand the role of the FAZ filament in *Leishmania*, we investigated the function of FAZ2 (LmxM.12.1120), which was identified as an essential FAZ filament protein in *T. brucei* (12). We generated a FAZ2 null mutant in *Leishmania mexicana* by sequential replacement of FAZ2 genes with antibiotic resistance markers. The integration of the resistance markers and the loss of the FAZ2 open reading frame were confirmed by PCR (Figure S1A). The parental cell line in which the deletions were performed expressed SMP1 endogenously tagged at its C-terminus with eGFP::Ty. SMP1 is an integral flagellum membrane protein and the tagged version acts as a marker for the flagellum membrane, enabling rapid analysis of changes to the flagellar pocket region of the *Leishmania* cell. We were readily able to generate the FAZ2 null mutant and the overall cell morphology was unaffected by the loss of FAZ2 (Figure 1A). However, the null mutant grew at a slower rate than the parental cell line with a slightly longer average doubling time in comparison the parental cells (7.1 (±0.6) hours vs 6.0 (±0.6) hours, t-test p=0.089) (Figure 1B). In the culture, we noticed the presence of cell ‘rosettes’ and cells connected to each other via their flagella (Figure 1C). To quantify this phenomenon, we analysed the cells directly from culture (Figure 1D). In the FAZ2 null mutant there was an increase in the percentage of cell rosettes (1.5% to 12%) and cells connected via their flagella (0% to 16.5%), with a concomitant drop in individual cells with one flagellum (82.5% to 59.5%). This suggests that there was a defect in cell segregation in the FAZ2 null mutant.

**Figure 1.**
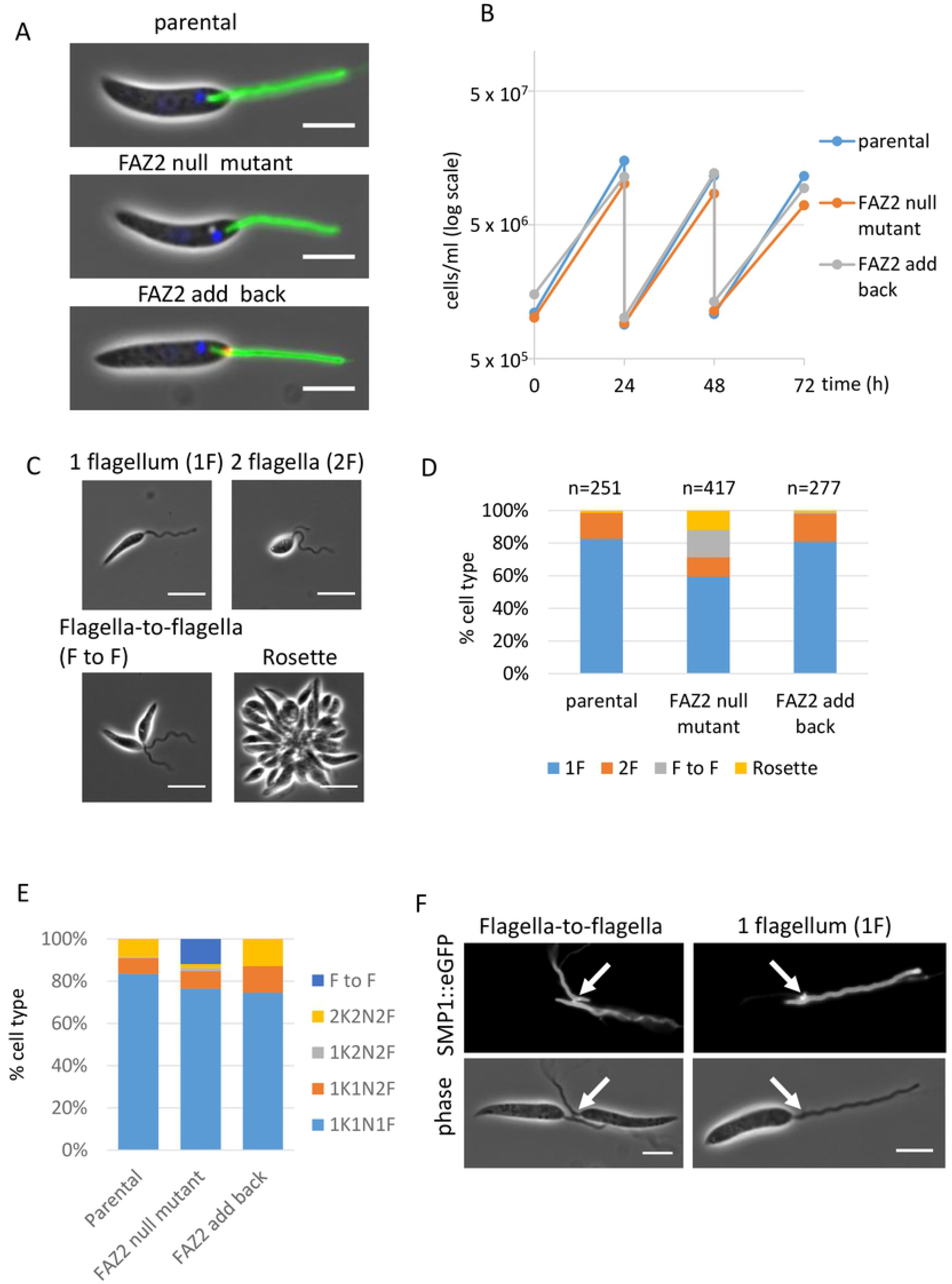
FAZ2 null mutant *Leishmania* parasites grew slower and had flagella-to-flagella connections. Light micrographs of parental, FAZ2 null mutant and FAZ2 add back cells expressing the flagellum membrane protein SMP1 tagged with eGFP::Ty (green) and the DNA is stained with Hoescht 33342 (blue). FAZ2 add back cells were expressing mChFP::FAZ2 (red). Scale bar is 5 μm. (B) Growth curve of the parental, FAZ2 null mutant and FAZ2 add back cells over a 72 h time period. (C) Light micrographs of cell types observed in culture. Scale bar is 5 μm. (D) Quantitation of cell types seen in culture for parental, FAZ2 null mutant and FAZ2 add back cells. (E) Cell cycle category counts for parental, FAZ2 null mutant and FAZ2 add back cells. F - flagellum, K - kinetoplast, N - nucleus, F to F - two cells connected via their flagella. (F) Light micrographs of FAZ2 null mutant cells expressing SMP1::eGFP::Ty showing the phase and SMP1::eGFP::Ty channels, with an example of two FAZ2 null mutant cells connected via their flagella and of a one flagellum FAZ2 null mutant cell that had a residual structure on the flagellum near the anterior cell tip (white arrow). Scale bar is 5 μm.

In *Leishmania* assembly of a new flagellum, and the duplication and segregation of the kinetoplast (mitochondrial DNA) and nucleus occur at set points during the cell cycle; therefore, the number of flagella, kinetoplasts and nuclei in a cell can be used to define its cell cycle stage. To assess whether there was a defect in cell segregation, we categorised the cells based on their number of flagella, kinetoplasts and nuclei (Figure 1E). FAZ2 null mutant populations had a reduced number of 1K1N1F cells, with the appearance of cells connected via their flagella. The drop in 1K1N1F cells and an increase in late stage cells (2K2N2F and cells connected by their flagella) suggests there was a problem in late stage resolution of cell division, which manifested in the phenotype of the unique flagellum-to-flagellum connection.

We saw many instances of two cells that were connected via their flagella with the connection point a short distance from the anterior tip of the cell body (Figure 1F, Supplementary movie 1, 2). When the flagella of connected cells were lying side-by-side at the base, a bridge was observed connecting the two flagella. In addition, on the flagellum of ~14% of 1F cells (n=65) a small SMP1 structure near the flagellum base; this structure was not observed on 1F parental cells (n=70) (Figure 1F). Given its location and presence only in FAZ2 null mutant cells, this SMP1-positive structure was likely to represent a remnant of the bridge that had previously connected two cells via their flagella.

To ensure that this flagellum-to-flagellum connection was a specific consequence of the loss of FAZ2 we introduced an ectopic copy of FAZ2 tagged with Ty::mChFP using an endogenous expression plasmid (23). Expression of the Ty::mChFP::FAZ2 ‘add-back’ protein was confirmed by western blotting, and the add-back protein localised to the expected position in the flagellar pocket neck by fluorescence microscopy (Figure 1A, S1B). The growth rate of the FAZ2 add-back cell line was similar to that of the parental cells and no cells connected by their flagella were observed, indicating the effects described above were indeed due to the loss of FAZ2 (Figure 1B, D, E).

### Cells lacking FAZ2 have a shorter flagellar pocket and a very short FAZ filament

Our previous work has shown that the deletion of FAZ5 led to changes in flagellar pocket shape (23). To determine if there was a change in the morphology of the flagellar pocket after FAZ2 deletion we measured the distance between the proximal end of the SMP1::eGFP::Ty signal and the flagellum exit point from the cell body (an estimate for flagellar pocket length) (Figure 2A). This distance was reduced in the FAZ2 null mutant in comparison with the parental and the FAZ2 add-back cells, suggesting that a shorter length of flagellum is housed within a flagellar pocket. To investigate the morphological changes to the flagellar pocket in greater detail we examined the anterior end of the cell by thin-section transmission electron microscopy (TEM) (Figure 2C, E). Thin-section TEM revealed that the overall organisation of the flagellar pocket was maintained, with both the bulbous region and the flagellar pocket neck region present. However, longitudinal images of the flagellar pocket revealed that both the length of the bulbous region (i.e. the distance between the basal body - i), and the length of the neck region (i.e. the distance between the flagellar pocket collar and between the flagellar pocket collar and the flagellum exit point - ii) were reduced in the null mutant (Figure 2C, D). This correlates with the reduced length of the flagellar pocket observed by light microscopy (Figure 2A, B).

**Figure 2.**
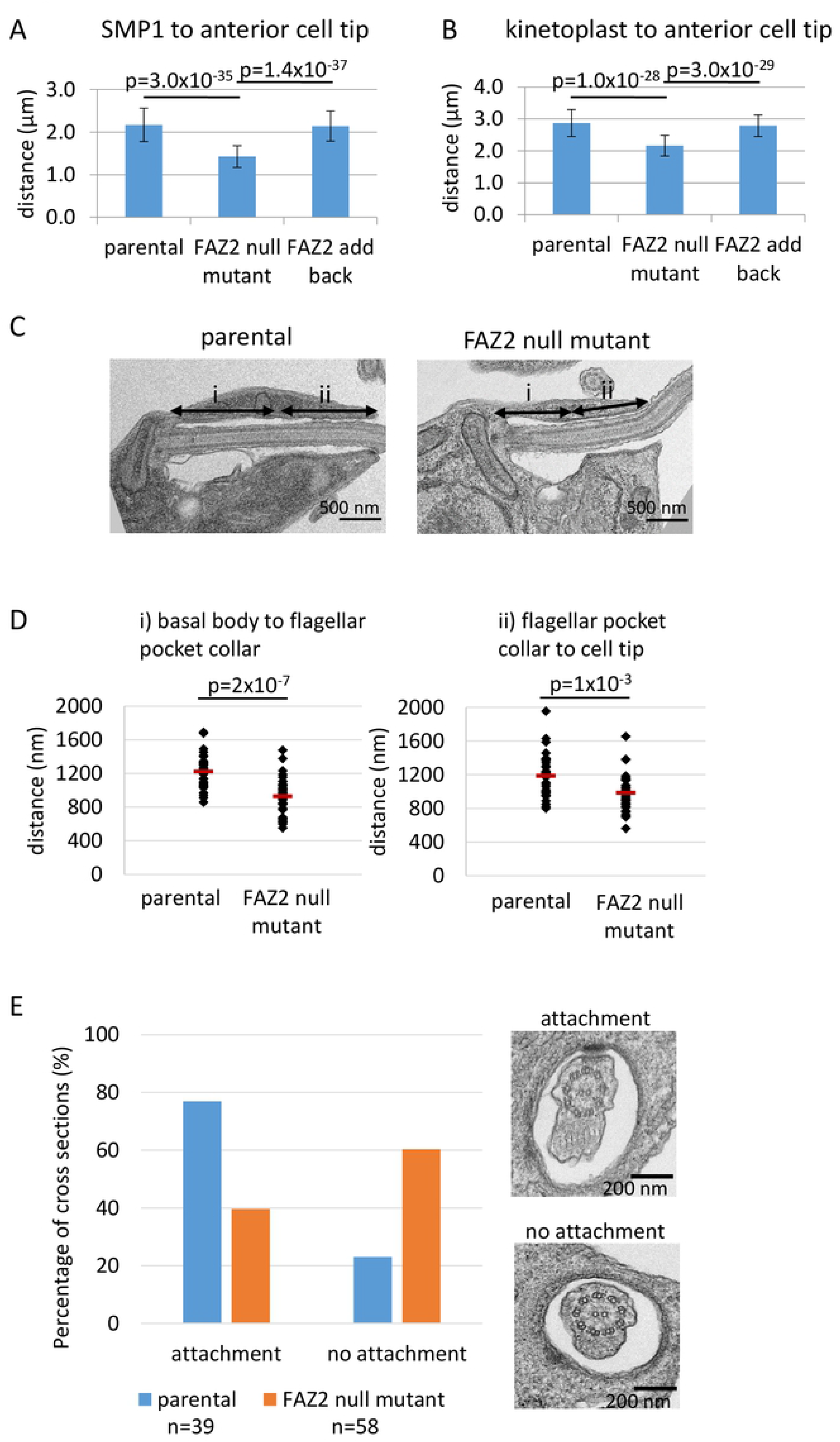
FAZ2 null mutants had an altered flagellar pocket shape and reduced level of flagellum attachment. (A) Measurement of flagellum length within the cell body as defined by the SMP1 signal body for the parental, FAZ2 null mutant and FAZ2 add back cells. 99 1K1N cells were measured for parental and FAZ2 add back cell line and 101 1K1N cells were measured for the FAZ2 null mutant, mean was plotted ±s.d. (B) Measurement of the distance between the kinetoplast and the anterior end of the cell body for the parental, FAZ2 null mutant and FAZ2 add back cells. 99 1K1N cells were measured for parental and FAZ2 add back cell line and 101 1K1N cells were measured for the FAZ2 null mutant, mean was plotted ±s.d. (C) Representative electron micrograph of longitudinal section through the flagellar pocket of a parental cell and FAZ2 null mutant cell. (i) represents the distance between the basal body and the flagellar pocket collar and (ii) represents the distance between the flagellar pocket collar and the anterior cell tip. Scale bar is 500 nm. (D) Measurement of i and ii highlighted in (C). Each measurement (parental n=34, FAZ2 null mutant n=35) was plotted with the mean represented as a red line. (E) Quantitation of the different cross sectional profiles across the flagellum and cell body observed as the flagellum extends through the flagellar pocket into the flagellar pocket neck region between the parental cells (n=39) and FAZ2 null mutant (n=58). Electron micrographs illustrate the two profiles based on the presence or absence of the attachment between the flagellum and the cell body. Scale bar is 200 nm.

The *T. brucei* FAZ2 homolog has a role in maintaining the attachment of the flagellum to the cell body (12). To examine if FAZ2 has a similar function in *Leishmania*, random TEM cross sections through the flagellar pocket were scored for the presence or absence of flagellum attachment to the cell body in both the parental and null mutant cells (Figure 2E). Deletion of FAZ2 led to a reduction in the number of cross sections in which attachment was observed; however, flagellum attachment to the cell body was still seen in nearly half the images examined. Next, we investigated whether FAZ2 deletion had overall effects on cell morphogenesis, by measuring the cell body length and width, and the flagellum length, in 1F1K1N cells of for parental, FAZ2 null mutant and FAZ2 add-back populations (Figure S1C-E). There were only minimal differences in these parameters between the different cell lines, showing that loss of FAZ2 did not have a great impact on overall cell morphogenesis.

To complement the thin-section TEM imaging we examined the flagellar pocket of the FAZ2 null mutant by serial electron tomography (Figure 3, Supplementary movie 3, 4), which allows detailed analysis of the FAZ cytoskeletal elements (2). Tomography data confirmed that the thin-section TEM images showing that the broad features of flagellar pocket organisation - the bulbous lumen and a flagellar pocket neck region demarcated by the collar were present in the null mutant (Figure 3D). Moreover, the microtubule quartet was observed in its normal position wrapping around the bulbous lumen of the flagellar pocket, passing through a gap in the flagellar pocket collar and extending into the flagellar pocket neck region (Figure 3D). However, the FAZ filament, which is normally found adjacent to the microtubule quartet in the flagellar pocket neck region, was much reduced in the FAZ2 null mutant, forming a short stub near the flagellar pocket collar (Figure 3D, F). In the parental cell the flagellum is connected to the cell body through a regular series of electron dense junctional complexes that form the primary attachment region (Figure 3A, B). In the equivalent position in the FAZ2 null mutant there were regions where the flagellar membrane appeared attached to the cell membrane, but these were not associated with the electron dense junctional complexes in the cell body (Figure 3D-F). An important feature of the *Leishmania* anterior cell tip is its asymmetry with the side associated with flagellum attachment extending a short distance further out along the flagellum than the surrounding cell body (Figure 3B). Interestingly, in the FAZ2 null mutant tomogram there was a ‘finger’ of cell body that extended along the flagellum, partially wrapping around the flagellum (Figure 3D, G, H). This extension did not contain cytoplasmic (sub-pellicular) microtubules, but contained electron dense structures reminiscent of the junctional complexes, associated with a region of attachment between the cell body and flagellum membrane. However, these densities were not arranged in the orderly pattern observed in the parental cell (Figure 3D, G, H).

**Figure 3.**
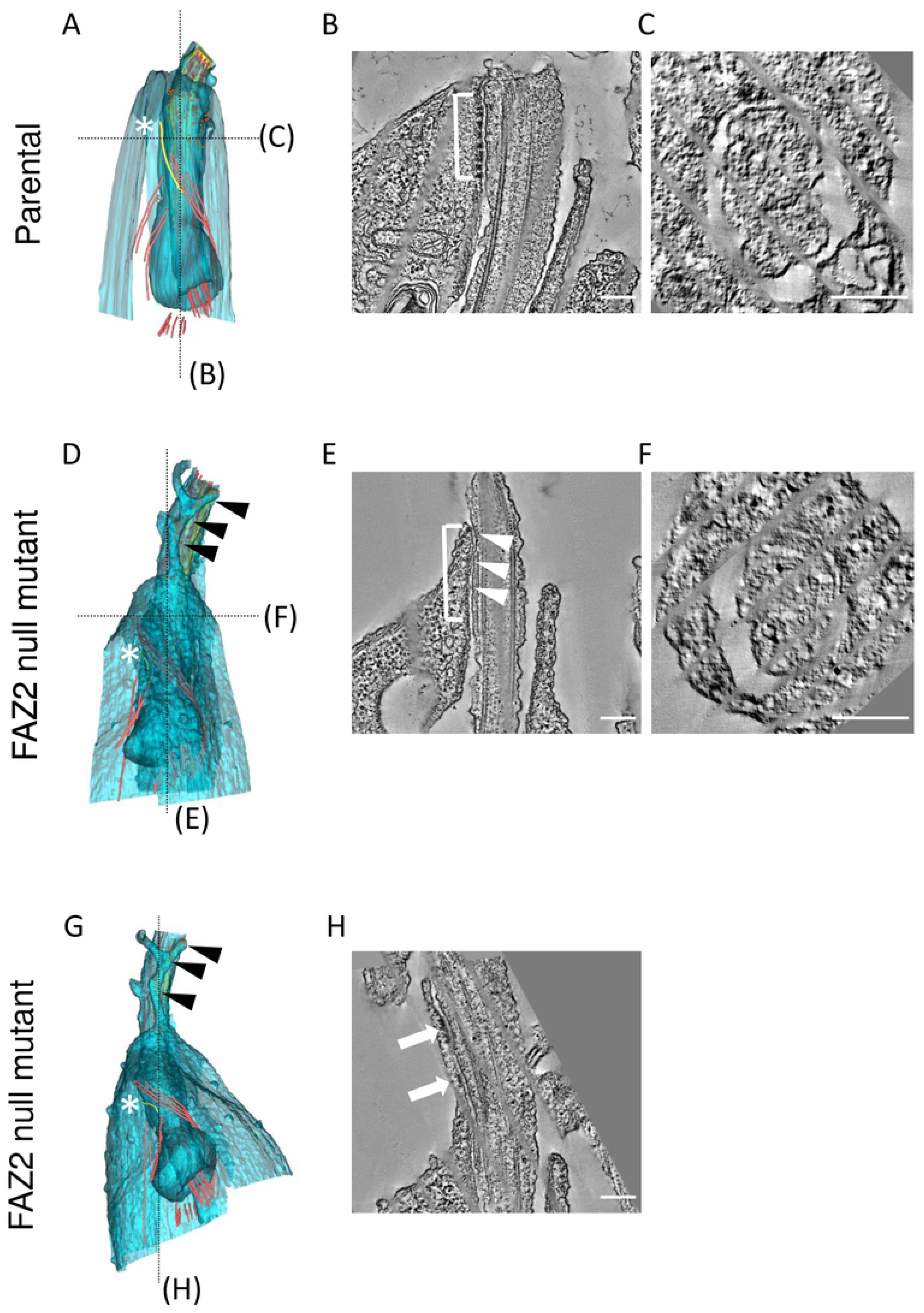
FAZ2 null mutants had a shorter FAZ filament and an extension of the anterior cell tip along the flagellum. (A) Model of flagellar pocket generated from tomogram of parental cell. FAZ filament in yellow marked by a white asterisk runs alongside the microtubule quartet in the flagellar pocket neck. Small orange spheres mark the junctional complexes attaching the flagellum to the cell body. Longitudinal tomogram slice through model in (A) along dotted black line. The regular electron dense junctional complexes mediate attachment of the flagellum (white bracket). The attached side of the cell body extends further along the flagellum than the opposite side. (C) Cross-sectional tomogram slice through model in (A) along dotted black line. White asterisk marks the FAZ filament. (D, G) Model of flagellar pocket generated from tomogram of FAZ2 null mutant. The FAZ filament in yellow is much shorter (white asterisk), whilst the microtubule quartet appeared unaffected by FAZ2 deletion. A projection of cell body extended along the flagellum. There were fewer junctional complexes (small orange spheres) with the majority found in the cell body projection (black arrowheads). (E, H) Longitudinal slices through models in (D, G) along the dotted black line. (F) Cross-sectional slice through model in (D) along the dotted black line. The flagellum was still connected to the cell body (E - white bracket) and distinct fibres connecting the cell body and flagellum membrane were seen (E - white arrowheads). Disorganised electron dense junctional complexes were also seen in the cell body extension of the FAZ2 null mutant (G - black arrowheads, H - white arrows). Scale bar is 200 nm.

### Deletion of FAZ2 disrupts the molecular organisation of the FAZ

The loss of FAZ2 resulted in errors in the morphogenesis of the anterior cell tip that appeared to affect the organisation and position of the FAZ. We therefore wanted to analyse the changes to the molecular structure of the major FAZ domains: i) the cell body, ii) the intermembrane and iii) the flagellum in more detail (4). We endogenously tagged FAZ proteins that represent these different domains with mChFP in the parental and FAZ2 null mutant cells and then imaged them with fluorescence microscopy (Figure 4). FAZ1 is found within the cell body FAZ domain, FAZ5 is located on the cell body side of the intermembrane domain, whereas FLA1BP is on the flagellum side of the intermembrane domain and ClpGM6 localises to the flagellum domain (2,9,11,21). In addition, FAZ10 was tagged as a marker of the flagellum exit point (2). There was no change in the localisation of FAZ10 between the parental and FAZ2 null mutant cells, with the FAZ10 signal forming a ring around the flagellum exit point from the cell body in both cell lines (Figure 4A). In the parental cells, the FAZ1 signal had the expected pattern with a ring around the flagellum and a short line parallel to the flagellum within the flagellar pocket neck; however, in the FAZ2 null mutant only the ring structure was observed, which was more pronounced than in the parental cells (Figure 4B). The loss of the short FAZ1 line signal correlates with the reduction in the length of the FAZ filament observed by cellular electron tomography.

**Figure 4.**
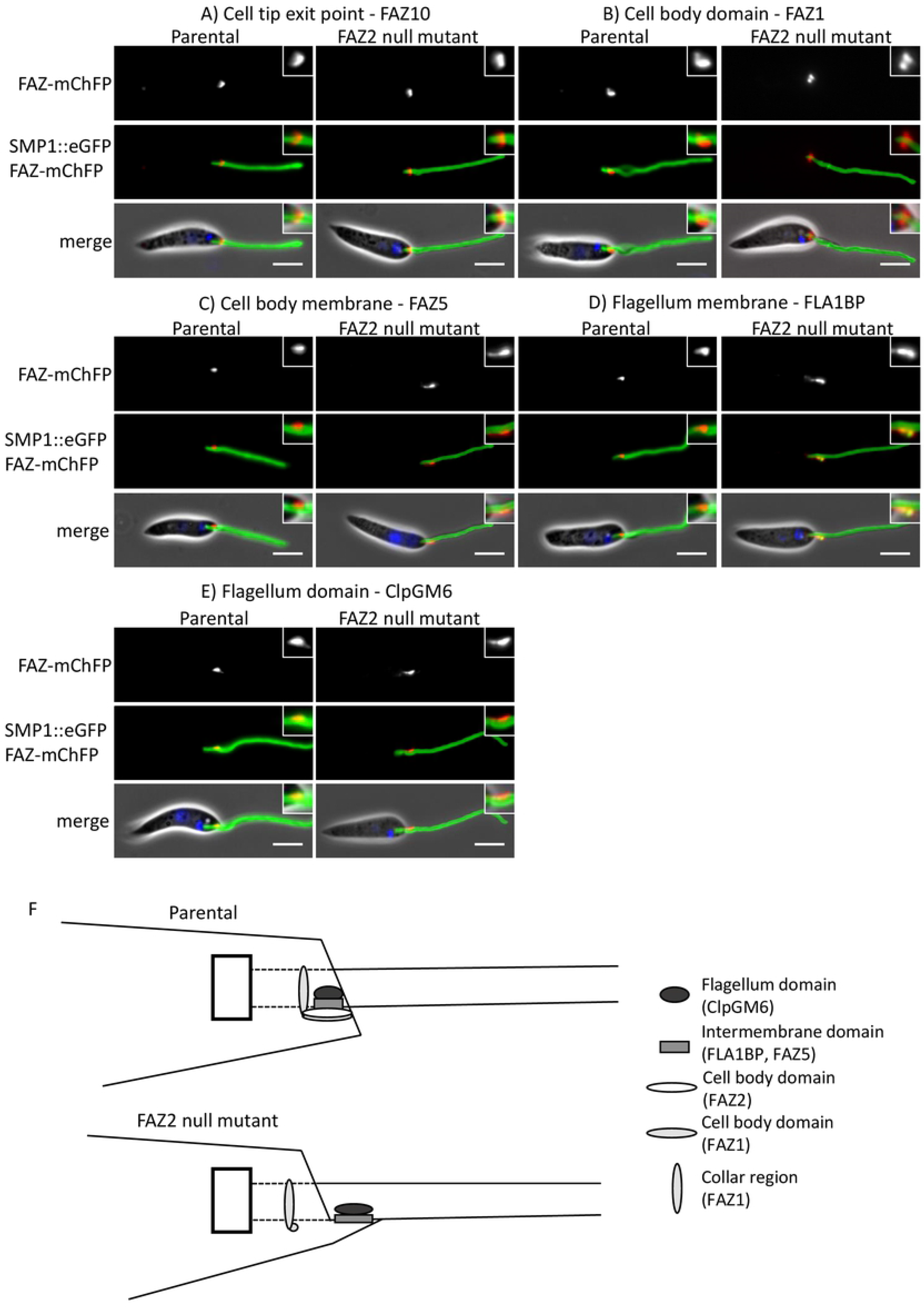
Deletion of FAZ2 affected the localisation of FAZ proteins in the FAZ membrane and flagellum domains but had little effect on FAZ proteins in the cell body domain. (A-E) Images of cells expressing FAZ proteins tagged with mChFP (red) representing the different FAZ domains in parental and FAZ2 null mutant cells. The flagellum membrane protein SMP1 is tagged with eGFP (green) and the DNA is stained with Hoescht 33342 (blue). The inset shows a zoomed in image of the FAZ protein localisation. Scale bar, 5 μm. F) Schematic of the FAZ domain organisation and anterior cell tip structure in the parental and FAZ2 null mutant cells.

FAZ5 signal was apparent as a short line parallel to the flagellum within the flagellar pocket neck region of parental cells, but in the FAZ2 null mutant the FAZ5 signal was not in the expected position; it appeared as a short line of signal alongside the proximal part of the flagellum, extending beyond the anterior cell tip (Figure 4C). This signal beyond the cell tip correlates with the extension of the cell body observed by electron microscopy (Figure 3D, G, H). FLA1BP and ClpGM6 had a similar localisation pattern to each other. In parental cells the FLA1BP and ClpGM6 signal appeared as a short line within the flagellum in the flagellar pocket neck region, which was asymmetrically positioned to one side of the flagellum (Figure 4D, E). However, in the FAZ2 null mutant cells FLA1BP and ClpGM6 were localised to a short region on one side of the flagellum with the strongest signal observed beyond the end of the cell body (Figure 4D, E). Together these data show that loss of FAZ2 resulted in changes to the organisation of the FAZ with the intermembrane and flagellum FAZ domains (FAZ5, FLA1BP, ClpGM6) disconnected from the cell body FAZ domain (FAZ1) and now localised outside the FAZ area in the neck region of the flagellar pocket, and potentially associated within an extension of the anterior cell tip found in the FAZ2 null mutant (Figure 4F). The mislocalisation of FAZ proteins in the FAZ2 null mutant provides evidence for a definitive role for the FAZ in late stage anterior cell morphogenesis.

### The flagellum-to flagellum connection in the FAZ2 null mutant are mediated by FAZ proteins

Next, we wanted to examine the structure of the flagellum-to-flagellum connection in detail. To do this we examined the localisation of the endogenously tagged FAZ proteins in cells that were connected via their flagella (Figure 5). In connected FAZ2 null mutant cells the localisation of FAZ10 and FAZ1 matched that observed in cells with unconnected flagella (Figure 5A, B). In cells expressing FAZ5 with their flagella connected, the most intense FAZ5 signal coincided with the connection and fainter signals were observed extending from the connection along the flagellum towards the cell body (Figure 5C). FLA1BP had a similar localisation to that of FAZ5, with the strongest signal coinciding with the connection between the flagella and fainter signals extending from this point along the flagella towards the cell bodies (Figure 5D). In cells with connected flagella ClpGM6 signal was present along the flagellum for ~2 μm from the anterior cell tip to the connection region. Unlike FAZ5 and FLA1BP, the ClpGM6 signal did not coincide with the structure mediating attachment between the two flagella and instead appeared within the flagellum directly adjacent to the connection (Figure 5E). This shows that the connection between the flagellum was associated with FAZ proteins from the intermembrane and flagellum FAZ domains.

**Figure 5.**
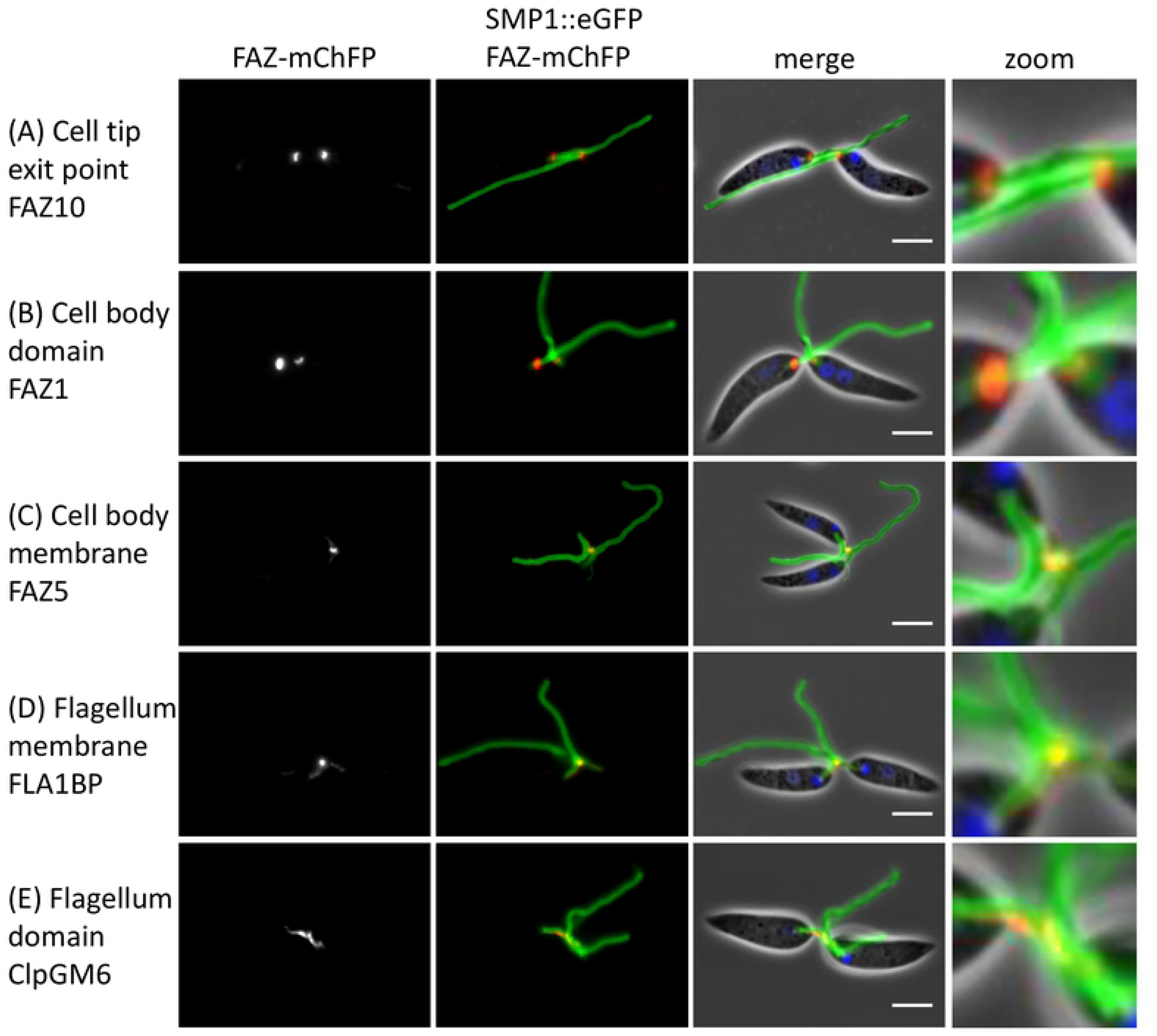
Flagella-to-flagella connections contained FAZ membrane domain proteins. (A-E) Images of FAZ2 null mutant cells expressing FAZ proteins tagged with mChFP (red) representing the different FAZ domains. The flagellum membrane protein SMP1 is tagged with eGFP (green) and the DNA is stained with Hoescht 33342 (blue). Scale bar, 5 μm.

We used scanning electron microscopy (SEM) to examine the flagellum-to-flagellum connection in the FAZ2 null mutant to determine at which point during the cell cycle it appeared (Figure 6A, B). The SEM micrographs showed that the two flagella were connected when the new flagellum was very short and had only just emerged from the flagellar pocket neck (Figure 6A, B). The connection persisted throughout the remainder of the cell cycle and for the majority of this time the flagellum-to-flagellum connection was not associated with the cell body. To examine the flagellum-to-flagellum connection further we used high-resolution SEM (Figure 6C) and serial electron tomography to generate a 3D reconstruction (Figure 6D, E, Supplementary movie 5). The high-resolution SEM micrographs showed that the connection between the flagella was not direct and was instead mediated by an additional membrane structure that acted as an intermediate. The electron tomography showed that in the intermediate connecting structure there were electron dense regions underlying the membrane that connected to each flagellum, which were associated with connecting fibres inside the flagella. These electron dense structures were highly reminiscent of the FAZ junctional complexes and correlate with the FAZ protein localisation, suggesting that the flagellum-to-flagellum connections between FAZ2 null mutant cells are orchestrated by FAZ proteins.

**Figure 6.**
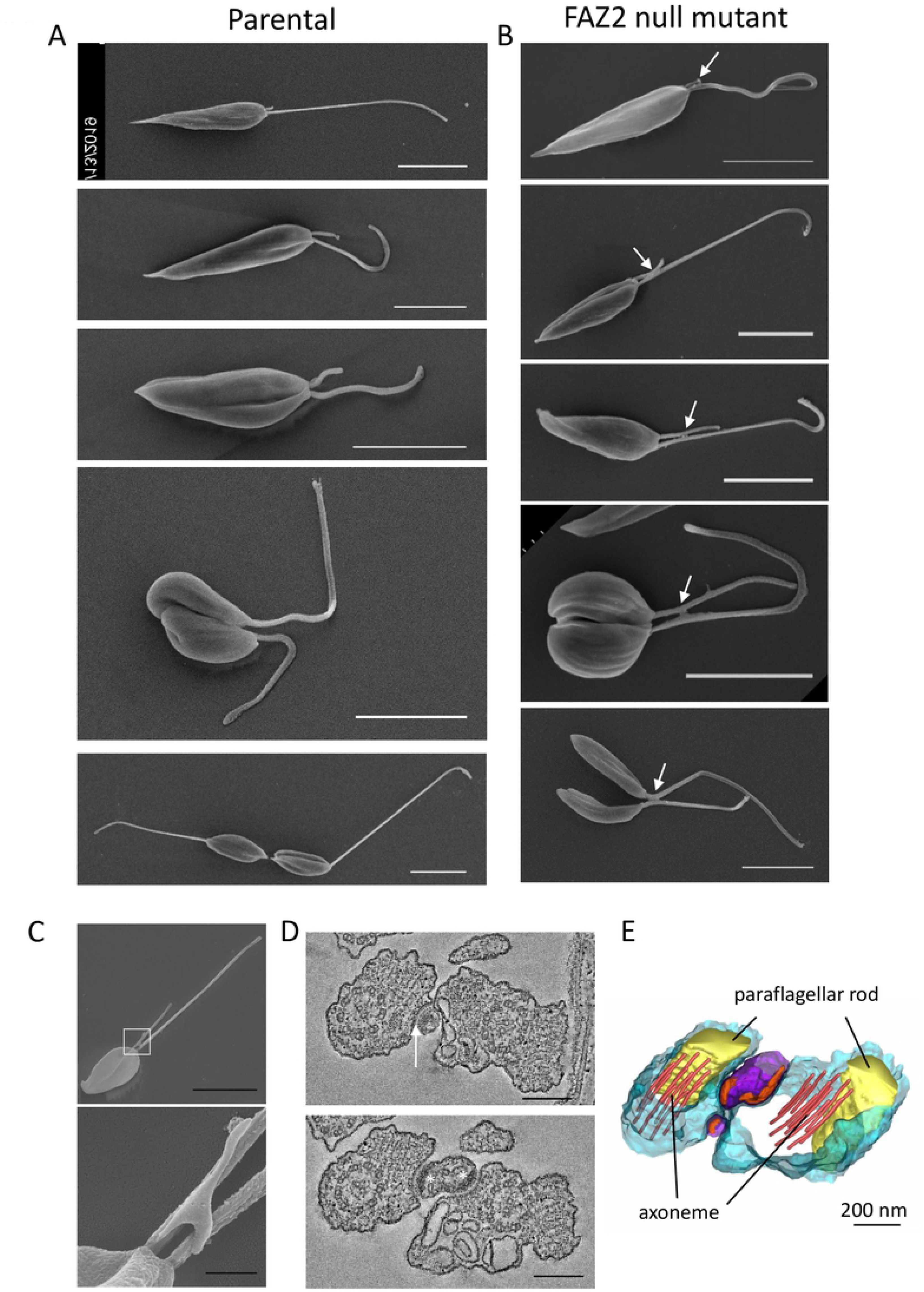
Flagella-to-flagella connections were mediated by a small membrane bound bridge structure. A, B) Conventional scanning electron micrographs of parental (A) and FAZ2 null mutant (B) cells through the cell cycle. The connection between is formed between the flagella (white arrow) as soon as the new flagellum exits the cell body. Scale bar is 5 μm. C) High-resolution scanning electron micrographs showing the bridge that mediates flagellum-to-flagellum connection (the white box indicates the position of the area enlarged beneath). Scale bar, 5 μm (A, B, C image on the left) and 500 nm (C, lower panel). D) Slices through the tomogram showing the connections (white arrow) between the flagellum and the connecting bridge structure. The asterisks indicate the electron density within the connecting structure. Scale bar, 200 nm. E) Model of the connection generated from tomogram of connected flagella in the FAZ2 null mutant.

### FAZ2 null mutant is unable to develop and proliferate in the sand fly vector

Given that the FAZ2 null mutant grew in culture we investigated whether the defect in anterior cell membrane resolution and the flagellum-to-flagellum connections affected the ability of the parasite to progress through its life cycle. We fed sand flies with blood containing either the parental, null mutant or add back cell line. We examined the development of the parasite by dissecting sand flies either 1-2 days or 6-8 days after the blood meal and categorised the parasite burden and its location in the sand fly gut (Figure 7, S2A). All three cell lines were able to establish an infection in the midgut 1-2 days after the blood meal, but the parasite burden in the null mutant was considerably lower than in the parental and add-back cell lines (Figure 7A). At 6-8 days after the blood meal very few sand flies were infected with the null mutant whereas for both the parental and add-back cells, the infection rate was above 70%, with more than 60% of sand flies having a heavy parasite burden (>1000 parasites/fly; Figure 7A). In the few sand flies in which the FAZ2 null mutant was present they were restricted to the midgut and had not colonised the stomodeal valve unlike the parental and add-back cells (Figure S2A). Together this showed that the deletion of FAZ2 decreased dramatically the proliferation and development of the parasite in the sand fly, suggesting that the defective in late stage resolution of cell segregation has stronger implications in the environment of the sand fly gut compared with the conditions of the in vitro culture. A potential alternative explanation for the inability of the FAZ2 null mutant to proliferate and develop in the sand fly is that the cells had a defect in motility (effective displacement) due to the flagellum-to-flagellum connection. To explore this idea, we examined the motility of the parental, null mutant and add-back cells by tracking the movement of 1000s of cells (Figure S2B). The cell tracks showed a clear reduction in processive movement in the FAZ2 null mutant in comparison with the parental and FAZ2 add-back cells, suggesting that the flagellum-to-flagellum connection impedes the ability of the parasite to move in a directional manner.

**Figure 7.**
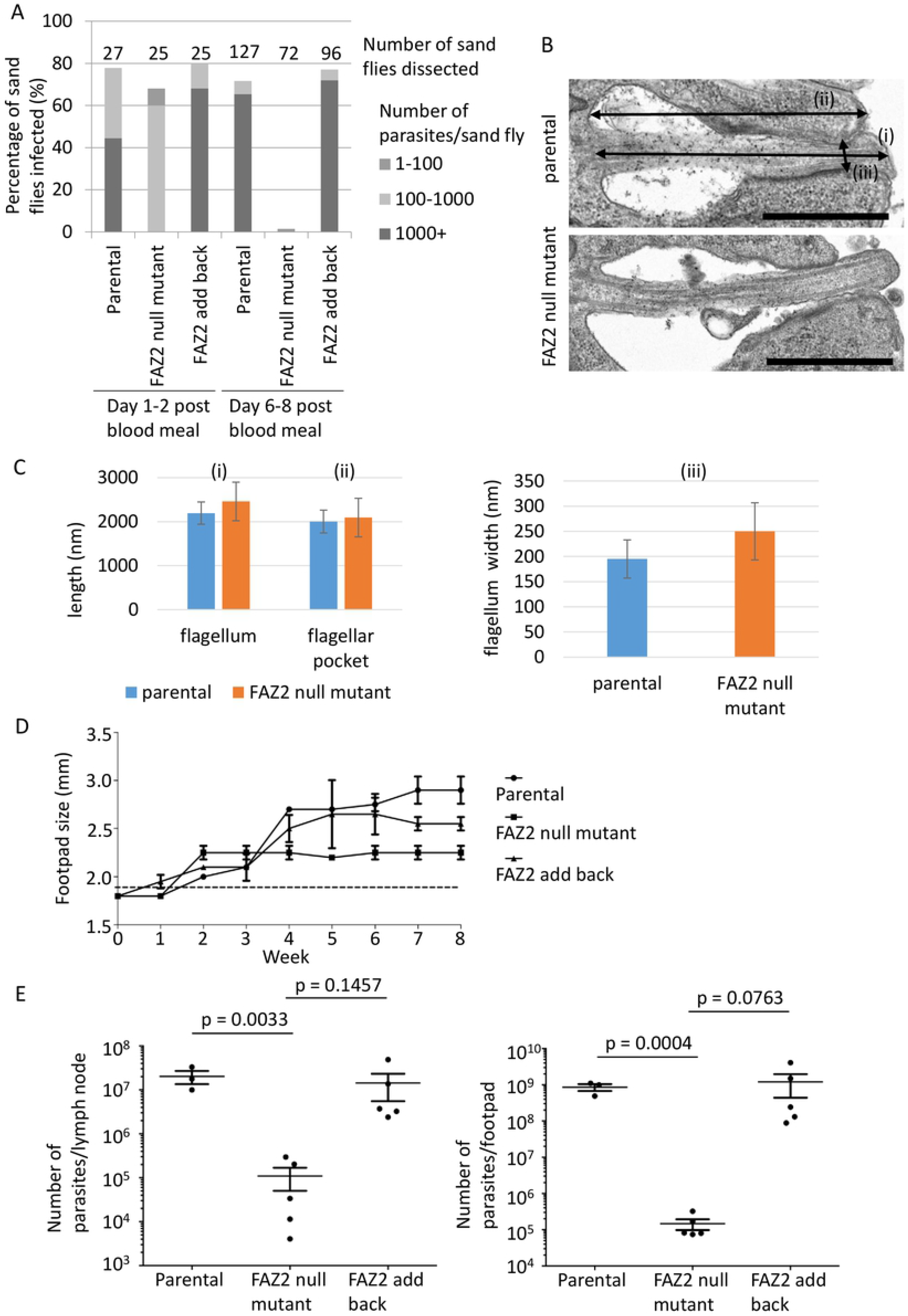
FAZ2 deletion severely affected the ability of *Leishmania* to proliferate and develop in the sand fly and dramatically reduced pathogenicity in the mouse. A) Analysis of sand fly infections using parental, FAZ2 null mutant and FAZ2 add-back cells. After 1-2 days post blood meal and 6-8 days post blood meal sand flies were dissected (number indicated above the column) and the parasite load in each fly was measured as heavy (1000+ parasites), moderate (100-1000 parasites), weak (1-100 parasites). B) Representative electron micrographs of longitudinal section through the flagellar pocket of a parental cell and FAZ2 null mutant axenic amastigotes. (i) represents flagellar pocket length, (ii) represents flagellum length, (iii) represents width of the flagellum at the constriction point. Scale bar is 1000 nm. C) Graphs for the 3 measurements in (B) with the mean plotted and error bars showing standard deviation for parental and FAZ2 null mutant axenic amastigotes. For flagellum length - parental cells n=16, FAZ2 null mutant n=24; for flagellar pocket length - parental cells n=16, FAZ2 null mutant n=27; for flagellum width - parental cells n=18, FAZ2 null mutant n=29. D) Measurement of mouse footpad lesion size during an 8-week infection time course with parental, FAZ2 null mutant and FAZ2 add-back cells. Error bars indicate standard deviation. E) Measurement of parasite burden at the end of the 8-week infection time course in the footpad lesion and the lymph node for the parental, FAZ2 null mutant and FAZ2 add back cells. Parasite number from each infection is plotted with the mean and the 95% SEM interval indicated, and p values were calculated using a two-tailed unpaired Student’s t-test (n=5).

### FAZ2 deletion affects amastigote structure and reduces pathogenicity in the mouse

In the mammalian host, *Leishmania* is an intracellular parasite that resides within a parasitophorous vacuole and has an amastigote morphology with a much shorter flagellum that only just extends beyond the cell body (24). During the promastigote to amastigote differentiation there is a reorganisation of the FAZ structure and changes in FAZ protein localisation (2). To understand whether FAZ2 deletion affected amastigote formation we triggered differentiation of promastigotes in vitro by reducing the pH and increasing the temperature (25). Over 72 hours the FAZ2 null mutant differentiated successfully and there were no apparent differences between the morphology of the null mutant amastigote and that of the parental and add-back amastigotes (Figure S3A).

To investigate the flagellar pocket region of amastigotes in greater detail we imaged these cells by thin-section TEM (Figure 7B). The null mutant flagellar pocket still had both the bulbous lumen and neck region. In longitudinal sections the flagellum appeared to extend beyond the cell body further (Figure 7B). Morphological analysis revealed that the flagellum of the FAZ2 null mutant amastigotes was on average slightly longer than that of parental amastigotes (2460±440 nm (n=24) vs 2194±252 nm (n=16), t-test p=0.02), with the mean length of the flagellar pocket essentially unchanged; however, the variations in flagellar/flagellar pocket length in the mutant cells was much greater (Figure 7C). Moreover, we noticed that the constriction point at the distal end of the flagellar pocket neck was wider in the FAZ2 null mutant than the parental cells (Figure 7C, 252±38 nm (n=29) vs 195±57 nm (n=18), t-test p=0.0003). This showed that deletion of FAZ2 affected flagellar pocket morphogenesis in the amastigote form.

Even though the FAZ2 null mutant was able to differentiate in vitro, we wanted to test its potential for growth in a mammalian host. We infected the footpads of mice with the parental, null mutant and add-back parasites and examined the infection progress over an 8-week time course (Figure 7D). Initially, infection with the FAZ2 null mutant resulted in an increase in footpad swelling over the first two weeks after infection, but beyond this point there was no further swelling, unlike that observed in parental and add-back infections. At the end of the time course the null mutant had caused much less swelling than the parental cells, with the add-back cells causing an intermediate level of swelling. Next, we assessed the parasite burdens in the footpad and lymph nodes (Figure 7E). There was a significantly reduced parasite burden in both the footpad and lymph nodes of those mice infected with the null mutant in comparison to the parental cells with the FAZ2 add back restoring the parasite numbers. The loss of FAZ2 had a great effect on the ability of the parasite to cause disease in the mouse, contrasting with the subtle phenotype observed in vitro.

A potential explanation for the reduced pathogenicity of the FAZ2 null mutant was that these cells were unable to establish an infection in phagocytic cells. To test this idea, we infected bone marrow derived macrophages with parental, null mutant and add-back cells for 2 hours and then followed the infection over 72 hours using stationary phase promastigotes (Figure S3B-D). The FAZ2 null mutant cells reached stationary phase with the same kinetics as the parental and add-back cells, though they had a lower final cell density. The infection indices (number of infected cells, number of parasites/cell) were similar between all the cell lines, indicating that the null mutant was still able to infect macrophages in vitro effectively (Figure S3C-D).

## Discussion

Accurate and successful cell division in *Leishmania* involves a new flagellum elongating alongside the old flagellum with the two flagella initially occupying the same flagellar pocket (26,27). The flagellar pocket then divides to generate two separate flagellar pockets each with its own flagellum. The cell membrane then ‘folds in’ forming a cytokinesis furrow that proceeds from anterior to posterior generating two daughter cells (26). The FAZ2 null mutant had a slightly reduced growth rate, with a small rise in cells in the late stages of the cell cycle, which was associated with defects in the morphogenesis of the anterior cell tip. This demonstrates an important role for FAZ2 and the FAZ filament in the resolution anterior cell tip membrane segregation during the late stages of cell division.

In parental cells the anterior cell tip around the flagellum exit site is asymmetric, with the side associated with the FAZ extending slightly further alongside the flagellum (2,28). We now demonstrate that this asymmetry is set by the FAZ, because deletion of FAZ2 causes an extension of the anterior cell tip, whilst deletion of FAZ5 results in the loss of the anterior cell tip asymmetry (Figure 3, Sunter *et al.*, 2019). In parental cells, the cell body FAZ domain and the intermembrane and flagellum FAZ domains are found in the same region along the length of the flagellum (2); however, in the FAZ2 null mutant these domains did not coincide, with the intermembrane and flagellum FAZ domain proteins being outside the FAZ region, distally separated from the FAZ filament remnant in the cell body. The exaggerated asymmetric extension containing intermembrane and flagellum FAZ domain components does not contain sub-pellicular microtubules, suggesting that it is ‘independent’ from the cell body proper. The simplest explanation for this phenotype is that it, in the absence of FAZ2, intermembrane and flagellum FAZ assemblage is ‘released’ from the physical link to the FAZ filament, and moves along the flagellum, within an anterior cell extension that elongates as the new flagellum extends.

The membranous bridge structure connecting the two flagella in dividing cells was present as soon as the new flagellum emerged from the flagellar pocket neck and, at this point, it was still connected to the cell body. As the new flagellum elongated the connection separated from the cell body. There is likely to be a limited length that the anterior cell tip can elongate to before the connection with the cell body breaks, resulting in the formation of the membranous structure connecting just the two flagella (Figure 8). This bridge structure was positive for SMP1, a flagellum membrane protein, suggesting that it originates from the flagellum or at least can receive flagellar membrane components. Within the flagellar pocket a boundary function likely operates to ensure that the flagellum membrane has a distinct protein complement (29); the loss of this boundary function could result in the movement of SMP1 to the connecting structure. However, SMP1 was not enriched on the cell body membrane so a loss of protein positional control seems unlikely. Our data, however, showed that this connection was likely formed from an extension of the anterior cell tip. In *T. brucei* transient interactions between flagella of different cells resulted in the transfer of fluorescent proteins between them (Imhof et al., 2016). The close apposition of the flagella and the connecting structure might result in the transfer of SMP1 by a similar mechanism.

**Figure 8.**
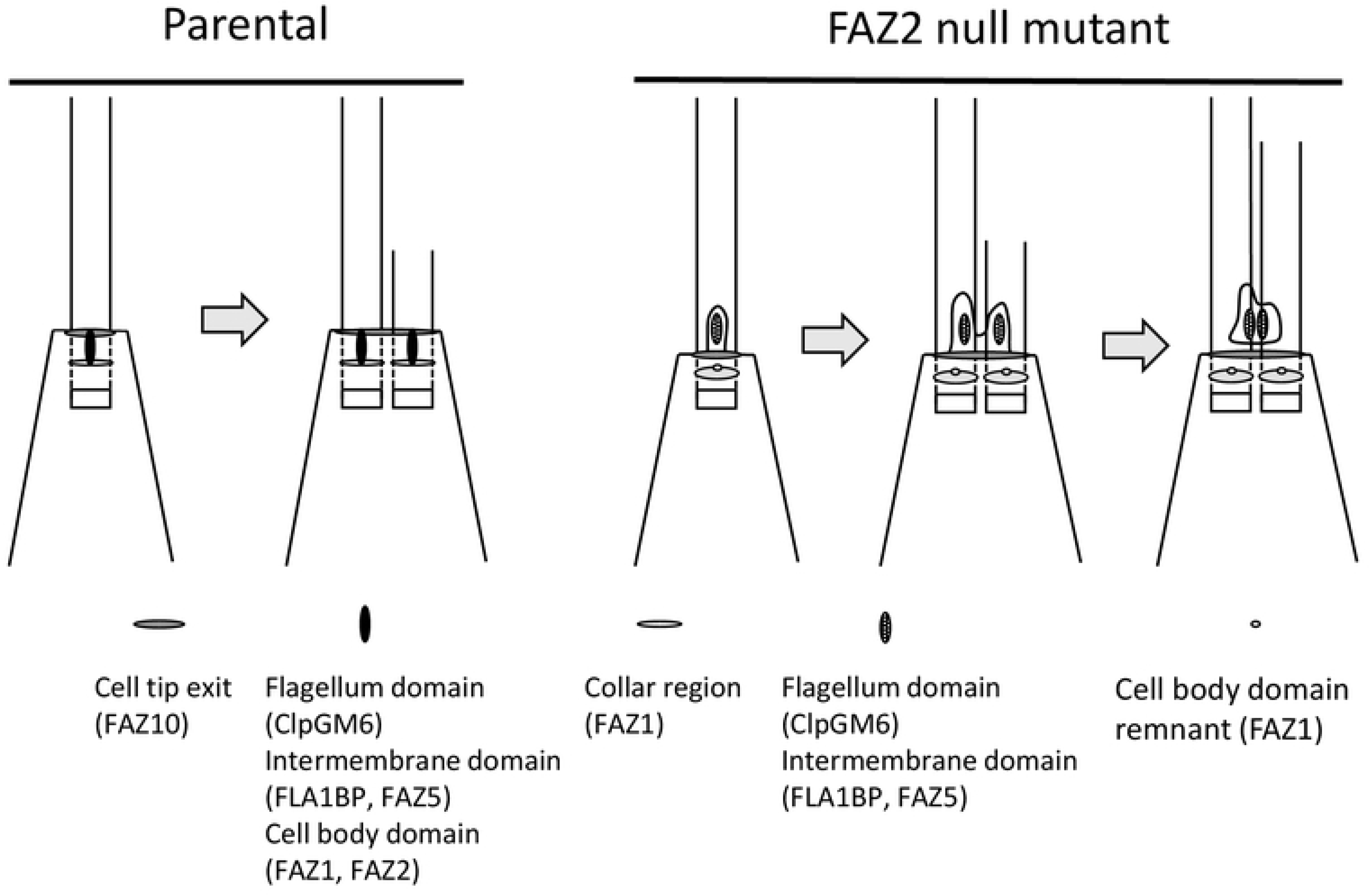
Model for the generation of the flagella-to-flagella connections. Plan view of the anterior end of the *Leishmania* promastigote, with the FAZ domains all vertically aligned: flagellum domain on top, next the intermembrane domain, then finally the cell body domain. In the parental cell the FAZ and flagellum assemble alongside the existing FAZ/flagellum structure. In the FAZ2 null mutant the flagellum and intermembrane FAZ domains are ‘released’ from their link to the FAZ cell body domain and move along the flagellum within an anterior cell tip extension. During new flagellum and FAZ assembly the newly assembled flagellum and intermembrane FAZ domains are also not linked to the cell body FAZ domain remnant and move along the new flagellum within an anterior cell tip extension that is connected to the existing cell tip extension along the old flagellum. As the new flagellum continues to grow xxthis extension separates from the cell body to leave the two flagella connected by a membranous bridge structure.

In the FAZ2 null mutant the membranous bridge between flagella is isolated from the cell body and, thus, unaffected by cell division resolution, which generates the unusual phenotype of post-division cells connected by their flagella. This suggests that, in *Leishmania* promastigotes, FAZ2 coordinates anterior cell tip separation and morphogenesis is FAZ2 dependent. In the absence of FAZ2, the anterior cell tip can divide, the collar can be segregated into two and two flagellar pockets formed and hence, cytoskeletal morphogenesis appears normal. However, membrane division (cell body and/or flagellum) is compromised, leading to alterations of asymmetry and the errors in the resolution of the anterior cell tip membrane.

The depletion of FAZ2 in *T. brucei* by RNAi had a catastrophic effect on the cell morphogenesis with the loss of flagellum attachment and destabilisation of FAZ proteins (Zhou et al., 2015). However, in *Leishmania* FAZ2 null mutants, flagellum attachment was reduced but not lost completely and although there were changes in FAZ protein localisation no large change in FAZ protein amount was observed. These differences are likely explained by the differences in the spatial organisation of the FAZ with the primary attachment zone in *Leishmania* not being as closely associated with the FAZ filament. This suggests a nuanced function of the *Leishmania* FAZ, with the FAZ filament playing a greater role in membrane resolution during cell segregation than actual flagellum attachment.

The FAZ2 null mutant parasites were able to survive within the blood meal surrounded by the peritrophic matrix in the midgut of the sand fly, albeit with a reduced infection density; however, these parasites were unable to proliferate and develop late-stage infections and hence did not colonise the cardia (i.e. the most anterior part of the thoracic midgut) and the stomodeal valve. A key step in the development of the parasite in the sand fly is the escape from the peritrophic matrix and attachment to the midgut epithelium with a recent study showing that motility is important for the parasite to complete its life cycle in the sand fly (30–32). We found that the motility of the FAZ2 null mutant in vitro was impaired likely due to the abnormalities in cell body-flagellum connections and the flagellum-to-flagellum connection impeding the ability of these cells to move effectively. This loss of directional movement could explain the lack of growth and development of this mutant in the sand fly. However, it is technically difficult to observe individual parasite movement in the sand fly, something that is common to all such studies and complicates the attribution of specific causal explanations for mutant behaviour. Here, as in other mutant studies, it is more likely that the reduced infection in the sand fly midgut is multifactorial, resulting from a combination of different phenomena.

The *Leishmania* amastigote flagellar pocket has a two-part structure with the bulbous lumen and the flagellar pocket neck region as found in the promastigote. However, the shortened amastigote flagellum only just extends beyond the anterior cell tip, and at the distal end of the flagellar pocket neck there is a constriction that squeezes tightly around the flagellum, coinciding with the localisation of FAZ2 (2). The loss of FAZ2 did not affect the ability of the *Leishmania* parasite to differentiate and the null mutant retained the two-part flagellar pocket organisation. However, the ultrastructure of the null mutant axenic amastigote flagellar pocket was more variable, with an increase in the width of the constriction point at the distal end of the flagellar pocket neck, showing that FAZ2 has an important role in maintaining amastigote flagellar pocket shape.

The FAZ2 null mutants were able to infect macrophages in vitro but had a reduced pathogenicity in vivo, with both a reduction in footpad swelling and parasite numbers recovered from the footpad and the lymph node. Thus, FAZ2 null mutant amastigotes struggled to survive and replicate over an extended period in the mouse. As with the promastigote in vivo phenotype there may be multiple effects leading to this result. However, changes to the amastigote flagellar pocket architecture in the null mutant may have contributed to the loss of pathogenicity in the mouse, because the increase in the width of the constriction point might well affect the ability of the parasite to control the exchange of material between the flagellar pocket and the extracellular milieu. The ‘relaxed’ constriction at the flagellar pocket exit might also increase parasite exposure to deleterious factors from the environment, hence reducing cell viability. The deletion of FAZ5 in *L. mexicana* altered the flagellar pocket architecture with the loss of the flagellar pocket neck region and these changes were associated with a large reduction in infectivity in the mouse (23). Together with our results here this suggests that flagellar pocket architecture is important for parasite pathogenicity in the mammalian host.

In summary, we have shown that the FAZ filament is critical for late stage morphogenetic resolution of the two nascent anterior ends of *Leishmania*. This clearly demonstrates the subtleties of the function of the FAZ not only as a crucial feature for flagellum attachment but also its role in membrane organisation at the anterior cell tip. This provides a deeper understanding of membrane-cytoskeletal interactions in the definition of individual cell form and the remodelling of that cell form at cytokinesis in this parasite.

## Materials and Methods

### Ethics statement

Experiments involving mice were conducted according to the Animals (Scientific Procedures) Act of 1986, United Kingdom, and had approval from the University of York Animal Welfare and Ethical Review Body (AWERB) committee.

### Cell culture

*L. mexicana* (WHO strain MNYC/BZ/1962/M379) promastigotes were grown at 28°C in M199 medium with 10% foetal calf serum, 40 mM HEPES-NaOH (pH 7.4), 26 mM NaHCO_3_ and 5 μg/ml haemin. Cells were maintained in logarithmic growth. Promastigotes were differentiated to axenic amastigotes by subculturing into Schneider’s Drosophila medium with 20% FCS and 25 mM MES-HCl (pH 5.5) at 34°C with 5% CO_2_, and grown for 72 h without subculture.

### Generation of FAZ2 deletion constructs, tagging constructs and FAZ2 add back construct

Deletion constructs were generated using fusion PCR as described (33). Regions comprising 500 bp of the 5’ UTR and 500 bp of the 3’ UTR of the *FAZ2* gene were combined with either the hygromycin resistance gene or the neomycin resistance gene by PCR to generate the deletion constructs. For tagging the corresponding ORFs and UTRs were cloned into pLEnTv2-YB plasmid (33). To produce the add-back cell line, the *FAZ2* gene was cloned into the XbaI and BamHI restriction sites of the constitutive expression plasmid described in (23). Constructs were transfected using a Nucleofector 2b as described previously (33).

### Light Microscopy

For live cell microscopy, cells were washed three times in PBS, resuspended in PBS with Hoescht 33342 (1 μg/ml) and then 5 μl of cell suspensions were placed on a glass slide. The cells were imaged using either a Leica DM5500B microscope with 100x objective and Neo 5.5 sCMOS camera or a Zeiss ImagerZ2 microscope with 63x or 100x objective and Hamamatsu Flash 4 camera. For cell swimming analysis, a 61 s video of 512 frames under darkfield illumination was captured using a 10x objective. Particle tracks were traced automatically as previously described (34).

### Transmission electron microscopy (TEM)

Cells were fixed in culture by the addition of glutaraldehyde for a final concentration of 2.5%. After 3 minutes, the cells were centrifuged (at 800g, for 5 min), washed in buffered fixative solution (0.1 M PIPES-NaOH buffer, pH 7.2, with 2.5% glutaraldehyde and 4% formaldehyde), resuspended in fresh buffered fixative solution and fixed overnight at 4°C. Cells were then washed five times in 0.1 M PIPES-NaOH buffer, pH 7.2 (including one 30-min wash in 50 mM glycine in 0.1 M PIPES-NaOH buffer), and post-fixed in 1% OsO_4_ in 0.1 M PIPES-NaOH buffer at 4°C, for 2h. Cells were washed five times in deionized water, then stained en bloc with 2% aqueous uranyl acetate overnight, at 4°C. Samples were then dehydrated in ethanol and embedded in Agar 100 resin. Thin-sections were stained with Reynolds’ lead citrate, before imaging on a Tecnai T12, equipped with a OneView 4×4 mega pixel camera (Gatan).

### Electron microscopy tomography

Ribbons containing serial sections of ~150 nm were produced from samples prepared for TEM as described above. Sections were stained with Reynolds’ lead citrate before imaging at 120 kV, on a Tecnai T12 with a OneView (Gatan) camera. Each individual tomogram was produced from a total of 240 4K × 4K pixel images (120 tilted images each of 0 and 90° axes, with 1° tilting between images) acquired automatically using SerialEM. Individual tomograms were produced using eTOMO (IMOD software package), and consecutive tomograms were then joined to produce serial tomogram volumes, using eTOMO. Tri-dimensional models from serial tomograms were produced by manual tracing and segmentation of selected structures using 3Dmod (IMOD software package).

### Scanning electron microscopy

Cells were fixed by adding glutaraldehyde to final concentration of 2.5% into the culture. Cells were harvested by centrifugation at 800 g for 5 minutes, the supernatant was removed and primary fixative was added (2.5% glutaraldehyde in 100 mM sodium phosphate buffer). After two hours, cells were washed twice in PBS and settled for 5 minutes onto round glass coverslips treated with poly-L-lysine. Coverslips were washed two times in PBS and the samples were then dehydrated using increasing concentrations of ethanol (30%, 50%, 70% and 90% v/v in distilled water, followed by three times in 100% ethanol). Samples were then critical point dried. Coverslips were mounted onto SEM stubs using silver DAG and coated with gold using a sputter coater. Images were taken on a Hitachi S-3400N scanning electron microscope at 5 kV with a 5.5 mm working distance. For high-resolution SEM, samples were imaged at 10 kV in a Zeiss Merlin Compact, using an in-lens SE detector and a 3 mm working distance.

### Sand fly infections

All parasites were cultivated at 23°C in M199 medium with 20% FCS, 1% BME vitamins, 2% sterile urine and 250 μg/ml amikin. Before infections parasites were washed three times in saline and resuspended in defibrinated heat-inactivated rabbit blood at 1 × 10^6^ promastigotes/ml. *Lutzomyia longipalpis* were maintained at 26°C and high humidity on 50% sucrose solution and 14 h light/10 h dark. Sand fly females, 3-5 days old, were fed through a chick skin membrane (35). Fully-engorged females were separated and maintained at 26°C with free access to 50% sucrose solution. They were dissected on days 1-2 and 6-8 post bloodmeal and the guts were checked for localisation and intensity of infection by light microscopy. Parasite loads were graded as described previously (36). Each cell line was used to infect sand flies in two independent experiments.

### Macrophage infections

Bone marrow derived macrophages (BMDMs) were grown in DMEM with 10% FCS and 10 ng/ml M-CSF at 37°C with 5% CO_2_. BMDMs were grown to confluence and then used to seed wells at 2.5×10^4^ cells/well. Promastigotes in log growth were split to 1×10^5^ cells/ml and grown to stationary phase over 5 days. The stationary phase promastigotes were used to infect the BMDMs for 2 h at a MOI of 5. After washing the cells to remove any free parasites the infected BMDMs were incubated at 34°C with 5% CO_2_ in DMEM for 3 days. At each time point BMDMs were fixed with methanol and the stained with the DRAQ5 and then imaged. Infected BMDMs and *Leishmania* parasites were then counted.

### Virulence assessment *in vivo* - footpad measurement and limiting dilution assays

All procedures were performed under Home Office Licence and in accordance with Institutionally-approved protocols. Strain virulence was assessed by footpad swelling and parasite burden (37). For experimental infections the parasites had previously been passaged through mice, isolated and transformed into promastigotes before being used. Groups of 5 female BALB/c mice (4-6 weeks) were infected subcutaneously at the left footpad using 2.0 × 10^6^ stationary promastigotes in 40 μl of sterile PBS. Infections were followed weekly by footpad measurement, and animals culled after 8 weeks using approved Schedule 1 methods prior to removal of footpad lesions and lymph nodes under sterile conditions. Samples were kept in M199 supplemented with 5 μg/ml gentamycin, and footpads digested with 4 mg/ml collagenase D for 2 h at 37°C. Lymph nodes and digested tissues were mechanically dissociated and filtered through a 70 μm cell strainer. Homogenates were resuspended in M199 supplemented with 20% FCS and serial dilutions (2-fold) performed in 96-well clear flat-bottom plates. Each sample dilution was performed in duplicate and distributed in at least three plates. Sealed-plates were incubated for 7-10 days at 25°C, wells visually analysed for the presence of parasites, and number of parasites calculated by multiplying by the dilution factors.

## Acknowledgements

We thank Dr Eva Gluenz (University of Oxford) for the kind gift of the *L. mexicana* SMP1::eGFP cell line, Dr Jessica Valli (University of Oxford) for help with the macrophage infection assays. This work was funded by a Nigel Groome studentship to CH, the Wellcome Trust to JCM (200807/Z/16/Z), MSMT (CZ.02.1.01/0.0/0.0/16_019/0000759) to PV and Research Center UNCE (204072) to KP. This work was initiated in the lab of Professor Keith Gull (University of Oxford), which is supported by the Wellcome Trust (104627/Z/14/Z). We would like to especially thank Keith Gull for his intellectual input through many conversations which helped to guide the development of this work and manuscript.

## Author contributions

J.D.S. designed research; C.H., J.D.S., R.Y., C.M.C.C.-P., F.M.-L., J.M., and K.P. performed research; C.H., R.Y., C.M.C.C.-P., F.M.-L., J.M., K.P., P.V., J.C.M., and J.D.S. analyzed data; and F.M.-L., P.V., J.C.M., and J.D.S wrote the paper.

---

Supplementary Figure 1. (A) Confirmation of FAZ2 gene deletion. gDNA from 4 null mutant clones and the parental cells was analysed by PCR. PCR confirmed that FAZ2 ORF was no longer present in the null mutant clones (1–3) and that the resistance markers had integrated correctly in clones (1–3). The neomycin resistance gene had not correctly integrated into clone 4 and this clone was discarded. FAZ2 null mutant clone 1 was used for all subsequent experiments. (B) Western blot confirming expression and expected size (174 kDa) of Ty::mChFP::FAZ2 using the BB2 antibody. The SMP1::eGFP::Ty and BB2 cross reacting band acted as a loading control. (C, D) Measurement of cell body length and width for parental (n=99), FAZ2 null mutant (n=101) and FAZ2 add back cells (n=99). Mean was plotted ± standard deviation for each measurement. (E) Measurement of flagellum length for parental (n=99), FAZ2 null mutant (n=102) and FAZ2 add back cells (n=97). Mean was plotted ± standard deviation for each measurement.

Supplementary Figure 2. (A) Migration of *Leishmania* in sand fly gut. Location of *Leishmania* parasites within infected sand flies at 1-2 and 6-8 days post blood meal. Stacked columns indicate the percentage of infected sand flies with parasites in various locations within the sand fly. FAZ2 null mutant was unable to migrate to the stomodeal valve. Percentage of infected flies for each cell line is indicated above each column. (B) Swimming tracks from videomicroscopy of parental, FAZ2 null mutant and FAZ2 add back cells. Scale bar is 50 μm.

Supplementary Figure 3. (A) Images of axenic amastigotes of parental, FAZ2 null mutant and FAZ2 add back cells expressing SMP1::eGFP::Ty. Scale bar is 5 μm. (B) *Leishmania* macrophage infections. Growth curve of parental, FAZ2 null mutant and FAZ2 add back cells to stationary phase - average of 3 replicates. (C, D) Proportion of infected macrophages and the number of *Leishmania* per infected macrophage at 0, 24, 48, 72 hours post infection - 0 h time point is after 2 hours of infection and removal of cells not taken up. For each time point between 487-1074 macrophages were analysed, n=3 for parental, FAZ2 null mutant and FAZ2 add back. Error bars show standard deviation.

Supplementary Movie 1. Movie of two *Leishmania* cells connected by their flagella.

Supplementary Movie 2. Movie of two *Leishmania* cells connected by their flagella.

Supplementary Movie 3. Movie of tomogram through parental flagellar pocket.

Supplementary Movie 4. Movie of tomogram through FAZ2 null mutant flagellar pocket.

Supplementary Movie 5. Movie of tomogram through bridge structure connecting two flagella.

